# From data to publication in a browser with BRC-Analytics: Evolutionary dynamics of coding overlaps in measles virus

**DOI:** 10.1101/2025.10.13.682095

**Authors:** Anton Nekrutenko, Danielle Callan, Marius Van Den Beek, Dannon Baker, David Rogers, Wolfgang Maier, Aysam Guerler, Hiram Clawson, Scott Cain, Kelsey Beavers, Michael Schatz, Maximilan Haeussler, Bjorn Gruning, Sergei Kosakowsky Pond

## Abstract

The analytical landscape of pathogen research is often fragmented, hindering transparency and reproducibility due to diverse genomic data sources, numerous software tools, and suboptimal integration methods. Here, we introduce BRC-analytics, a novel browser-based environment that unifies authoritative sources of genomic data with community-curated best analysis practices on a freely accessible public computational infrastructure. We demonstrate its capabilities by analyzing the evolutionary dynamics within the P/V/C locus of the measles virus, a complex system involving overlapping coding regions and RNA editing. Our analysis, conducted entirely within BRC-analytics, reveals asymmetric evolution of the locus’s reading frames under distinct selective pressures. BRC-analytics streamlines the entire research process—from data collection and primary analysis (e.g., variant calling) to interpretation (e.g., using integrated JupyterLite notebooks and LLMs) and publication—into a single web browser session. This eliminates the need for local installations and manual data transfers, implicitly tracking provenance and ensuring reproducibility. The platform’s goal is to provide true data-to-publication functionality, making advanced pathogen genomics accessible to a broader research community, regardless of their computational expertise or access to infrastructure.

## Introduction

Every research study, from data collection to publication, involves four fundamental steps:

1. **Data Collection**: Necessary datasets, including reference genomes, annotations, and sequencing reads, are typically found in standard repositories like NCBI or EBI. However, these datasets often require downloading before use.
2. **Primary Analysis**: This stage involves processing sequencing reads to generate outputs such as variant lists and transcript counts, using various standard tools and pipelines. A significant challenge for many researchers is determining the appropriate tool combinations for their data and experimental design, as well as the complexities of installation, configuration, and execution environments. While LLM chatbots can assist with installation and configuration, they do not inherently improve analysis reproducibility. Furthermore, while modern laptops and desktops can carry out many analyses locally, large-scale studies with thousands of samples demand substantial computational resources.
3. **Interpretation**: This phase often lacks dedicated tools, relying instead on general utilities like spreadsheets or custom code in computational notebooks. A major drawback is that custom code is rarely versioned or documented, making these analyses difficult for external researchers to reproduce.
4. **Publication**: The final step where findings are disseminated.

BRC-analytics uniquely integrates all these steps into a single web browser session, streamlining the entire research process. It directly accesses reference genomes and annotations as well as all publicly available data from the sequence read archive (SRA) and combines it with community-curated best-practice analysis workflows and tools. BRC-analytics runs on free public computational resources, requiring no additional investment from the user.

To highlight the functionality of BRC-analytics, we picked the task of genomic pathogen surveillance. In this analysis, data from multiple spatially and temporally distributed samples are processed to identify and classify sequence variants. This is done by aligning reads against a reference genome, calling variants, and generating variant lists: locations and identities of nucleotide changes found in each sample. These variants are further processed to uncover underlying patterns such as dynamics of variants in space and time, and ultimately estimate their potential effects on the fitness of the pathogen. Before moving to practicalities, let us deconstruct this analysis into data collection, primary analysis, interpretation, and publication strategies mentioned earlier (Table 1).

**Table 1.**
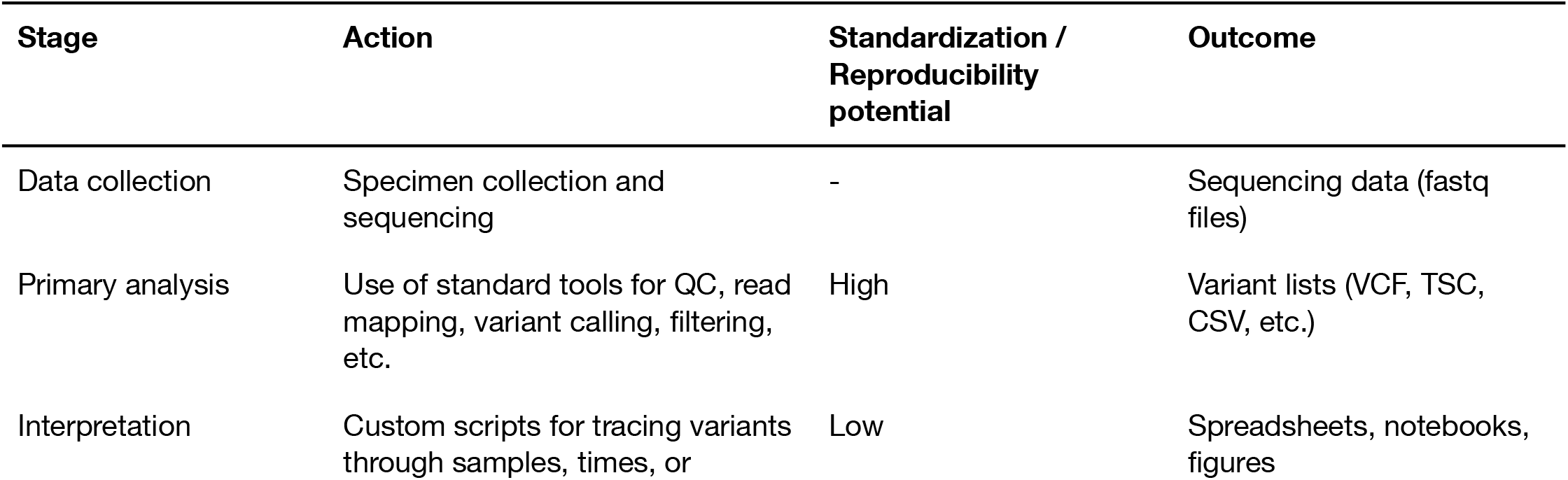
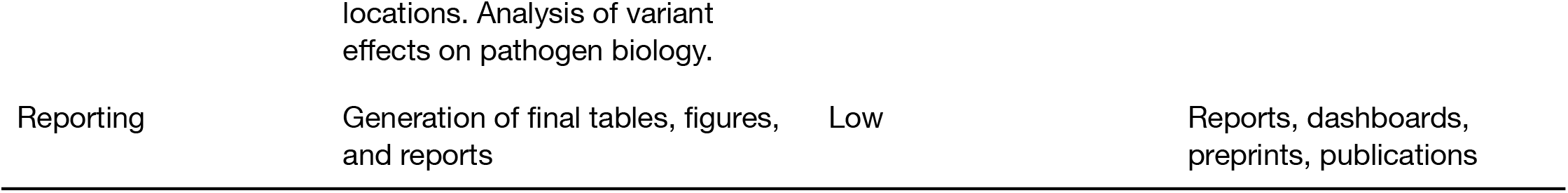
The four stages of a biomedical research project: a pathogen surveillance example.

Outside of BRC-analytics, all of these steps will usually be performed separately. Suppose a student in a lab performs a re-analysis of published surveillance datasets. These datasets—fastq files from the sequence read archive (SRA) such as NCBI or EBI—are first downloaded to local computers. Next, these data need to be mapped against the suitable reference genome for the pathogen in question. For this, the reference also needs to be located and downloaded. At this point, using a combination of tools or workflows, read data are reduced to a list of variants. Depending on how (individual tool, or Snakemake, Nextflow, or Galaxy workflows) and where (one’s laptop, an institutional cluster, a cloud, a Galaxy instance) this is done, the results may be more or less reproducible. The interpretation stage involves filtering, reshaping, and summarizing the list of variants. It is done using “analytical frameworks” such as spreadsheet applications or notebooks (Jupyter, RStudio). This is the least reproducible stage of the analysis because spreadsheet formulas and/or custom code created in notebooks are rarely versioned or documented. The entire process is usually condensed into a version of “…*analyzed using a collection of custom Python scripts*” found in tens of thousands of manuscripts. Because BRC-analytics integrates all these steps into a single environment, it automatically tracks provenance, saves all parameters, intermediate datasets, and code snippets in notebooks, thereby addressing most reproducibility concerns.

As a biological system for highlighting BRC-analytics functionality we selected the measles virus (MeV) for two reasons. First, there are a number of recent surveillance datasets available for this pathogen. These datasets are not very large, representative of a typical small-to-medium research lab. Such labs comprise the majority of single PI groups in the US (based on the analysis of R01-equivalent awards for 2024 found at [1]. RO1-equivalent are DP1, DP2, DP5, R01, R37, R56, RF1, RL1, U01, R35). Thus we can show how

BRC-analytics benefits the majority of biomedical researchers. Second, the MeV genome encodes the P/V/C-locus with two sets of overlapping reading frames (Fig. 1). Analysis of genetic variants within overlapping coding regions is challenging in practice, and allows us to highlight *impromptu* analytical capabilities of the BRC-analytics environment. A mutation occurring in protein-coding regions changes the nucleotide sequence of underlying codons. Due to redundancy of the genetic code such change may or may not alter the amino acid encoded by the codon. If a mutation does change amino acid, such mutation is called non-synonymous. Conversely, a mutation that does not lead to amino acid change is called synonymous. But what if a given gene codes for multiple proteins via overlapping reading frames (Fig. 2)? In this case the concept of “synonymous” or “non-synonymous” becomes relative to a frame [2,3]. For example, due to the organization of the standard genetic code most changes in the third codon position are synonymous.

**Figure 1.**
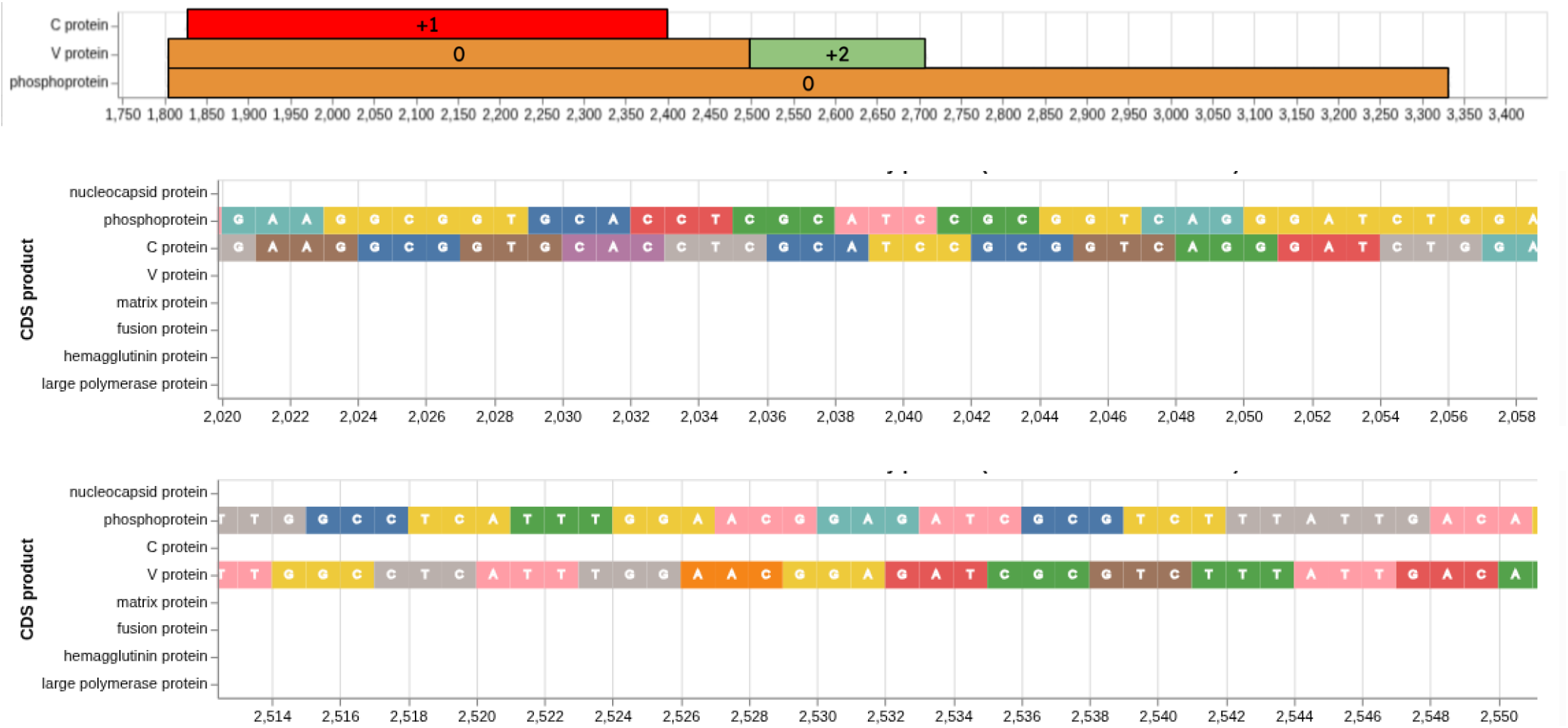
Schematic representation of overlapping reading frames within the P/V/C locus of measles virus, *MeV*, (upper pane). C (middle pane) and V (lower pane) are shifted +1 and +2 relative to the P frame, respectively. The P protein is translated from the primary mRNA transcript, while the V and C proteins are produced from the same locus through different mechanisms. The C protein, a small basic protein, is expressed by an alternative translational initiation mechanism. It is translated from an overlapping open reading frame (in +1 phase relative to P) that begins 19 nucleotides downstream of the P/V start codon. This strategy allows for the independent expression of the C protein from the same mRNA transcript that codes for P and V. The V protein is produced through cotranscriptional RNA editing. This process involves the viral polymerase inserting a non-templated G residue at a specific site within the P/V mRNA transcript. This pseudo-templated insertion triggers a +2 ribosomal frameshift, leading to the translation of a new V ORF. The mechanism, often referred to as “polymerase stuttering,” occurs when the viral RNA-dependent RNA-polymerase repeatedly reads a single template cytosine within a short G run that is part of a larger polypurine tract.

**Figure 2.**
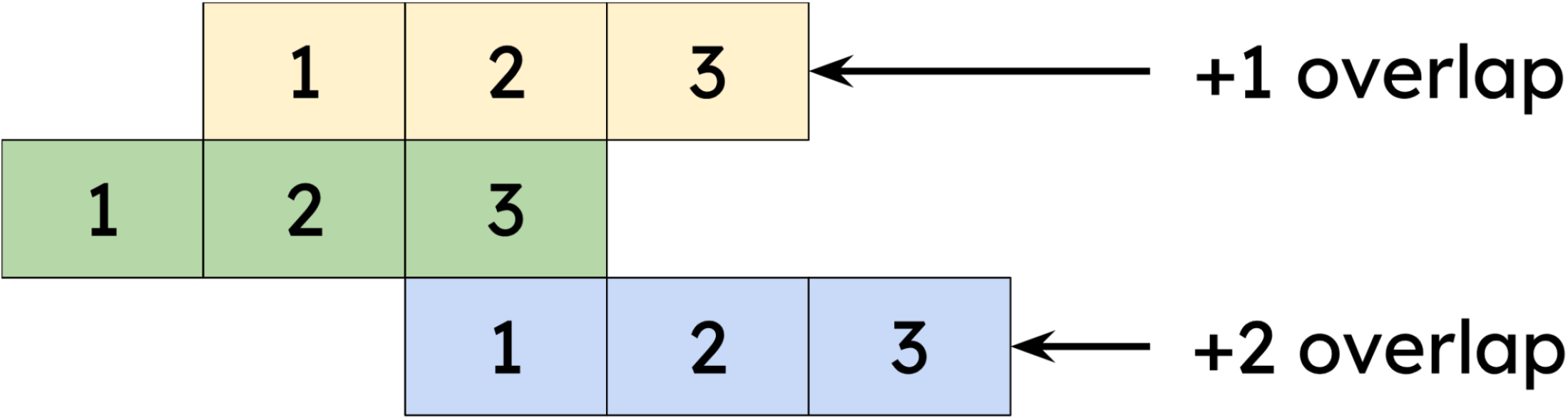
Correspondence of codon positions in +1 and +2 overlaps

However, a third codon position corresponds to the second codon position in +1 overlap and any change in the second codon position is always nonsynonymous. In other words a synonymous change in one frame will most often be non-synonymous in the other creating interesting scenarios for the evolution of proteins encoded by overlapping reading frames. Overlapping reading frames are rather common in viral genomes [4], and the challenges inherent in their analyses are well-recognized; in many cases such dual (or triple)-coding regions are excluded altogether. For instance, HIV-1, Influenza A virus, SARS-CoV-2, Hepatitis C virus, and norovirus all contain one or more overlapping reading frames.

In this manuscript we demonstrate how a routine surveillance analysis can be performed entirely using the BRC-analytics environment. We begin by obtaining all necessary data and applying a “best-practice” variant analysis workflow to create the initials list of variants. We then demonstrate how BRC-analytics allows custom analyses using a built-in JupyterLite functionality to perform non-standard analyses of overlapping coding regions and tracing evolutionary dynamics of nucleotide substitutions. This approach ensures transparency and reproducibility by implicitly tracking provenance, ultimately enabling researchers to conduct sophisticated evolutionary and functional analyses without local installations or manual data transfers.

## Results and Discussion

### What is BRC-analytics?

BRC-Analytics is an online, browser-based analysis environment designed to make comprehensive and reproducible genomic analyses of infectious diseases accessible to everyone. Developed under the

NIAID-funded Bioinformatics Resource Centers (BRCs) program, it leverages the Galaxy workflow system. The key features of the platform are listed below and also shown in Figure 3.

**Figure 3.**
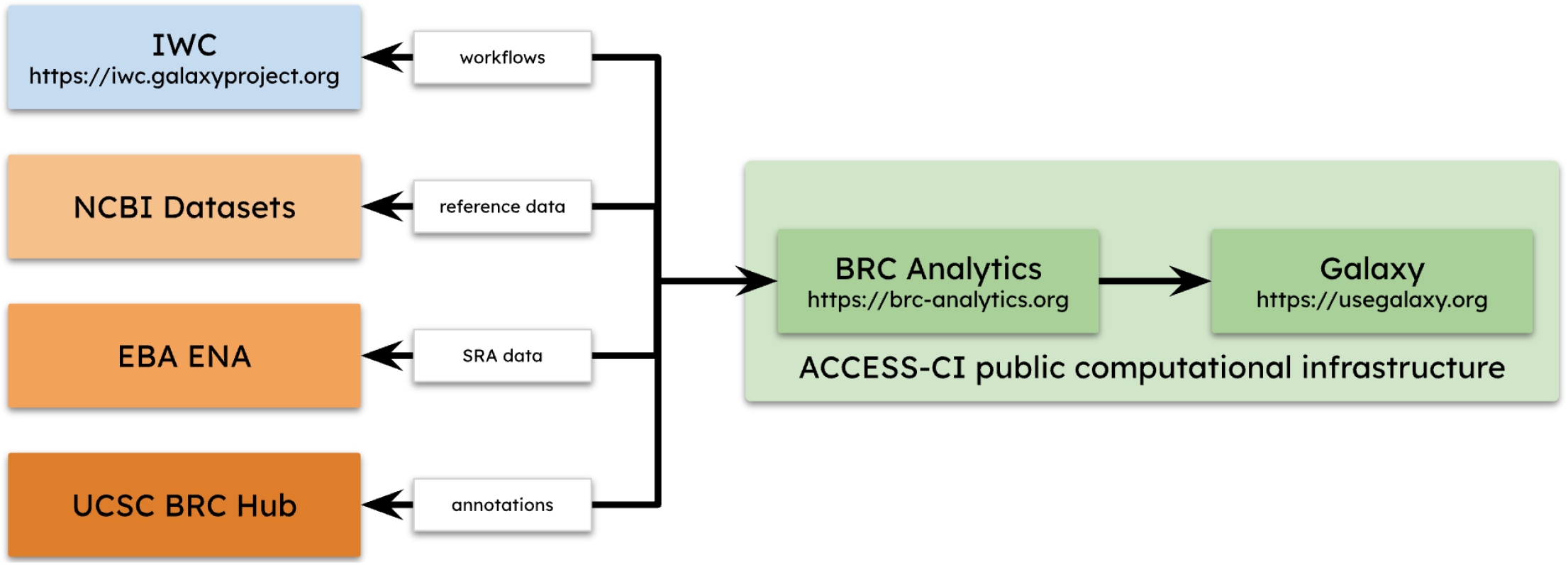
Main components of BRC-analytics.

#### Streamlined Research Process

Users can begin with raw sequencing reads and achieve publication-ready results without the need for local software installations or manual data transfers between tools.

#### Integrated Data

BRC-Analytics combines authoritative genomic data from NCBI Datasets with community-curated best-practice analysis workflows. These workflows cover essential steps like quality control, read mapping, variant identification, and annotation. Integrated data sources include <u>NCBI Datasets:</u> Populates BRC-Analytics with a list of organisms and assemblies, serving as the primary source for reference data. Currently, it lists 1,601 genome assemblies for 1,149 distinct pathogen, host, and vector taxa, with plans for continuous expansion. <u>UCSC Genome Browser:</u> Provides genome annotations (gene coordinates, regulatory elements, etc.). This integration allows for flexibility in including both reference and community annotations. <u>EBI ENA:</u> Facilitates access to public sequence read archive (SRA) data through local caching of SRA metadata for quick searches and an ENA API for read-level data.

#### State-of-the-art Analysis Workflows

These workflows cover essential steps like quality control, read mapping, variant identification, and annotation. BRC-Analytics uses Galaxy for launching and running workflows and individual analysis tools. Galaxy also serves as an environment for interpretive analyses, supporting interactive tools like Jupyter, RStudio, and JupyterLite.

#### Cloud-Based and Reproducible

The platform utilizes cloud-based computation, versioned workflows, and interactive visualizations to create a seamless interface. This approach unifies data and analytical capabilities, making advanced pathogen genomics available to a wider research community.

#### Computational Infrastructure

The substantial computational and storage resources required for BRC-Analytics are provided by the ACCESS-CI infrastructure in the US. BRC-Analytics and Galaxy are hosted on servers at the Texas Advanced Computing Center (TACC), a key component of ACCESS-CI.

### Selecting genome: Using NCBI Datasets for reference data (Video 1)

The analysis begins at https://brc-analytics.org where we search for *Morbillivirus hominis*—a taxonomic name for measles virus (a new standard for official scientific names implemented by the international committee on taxonomy of viruses; https://ictv.global/). The search yields a single assembly, GCF_000854845.1, corresponding to the only RefSeq genome for measles. In the remainder of this document we refer to the measles virus as MeV.

### Replicating this analysis

Everything described in this manuscript can be re-examined and replicated. The table below provides links and description to key artifacts of this analysis:

Clicking the history link will allow you to import that history in your own workspace and re-examine or re-run every step.

### Selecting analysis: Using community-curated workflows (Video 1)

Our goal is to perform a variant identification analysis: sequence data (reads) are mapped against a reference genome and mismatches between the reads and the genome are evaluated to identify likely changes. This procedure requires multiple tools for evaluation of data quality, read mapping, and variant identification along with auxiliary utilities. A standard and robust way to run a sequence of analyses like this is to assemble tools into a workflow. There are many community developed and maintained collections of workflows covering a very broad scope of common sequence analysis scenarios. To assist users with selecting a suitable workflow, BRC-Analytics will try to match an assembly to a list of applicable candidates. For the MeV reference genome, BRC-analytics suggests two variant calling workflows. For this study we are using a workflow called “*Variant calling and consensus construction from paired end short read data of non-segmented viral genomes*” (Fig. 4). This workflow was specifically designed for non-segmented viruses with a low-to-medium evolutionary rate such as MeV. Detailed information about the workflow can be found at [IWC](https://iwc.galaxyproject.org/workflow/generic-non-segmented-viral-variant-calling-main/)—Galaxy’s workflow repository and at [Dockstore](https://dockstore.org/workflows/github.com/iwc-workflows/generic-non-segmented-viral-variant-calling/main:main?tab=info). The workflow consumes samples (deep sequencing short read datasets, representing a viral population in a single host), and produces various outputs: a summary of variants found in each sample, consensus sequence for each sample as well as auxiliary outputs such as QC summary for all samples (see *Making sense of variants* section below).

**Figure 4.**
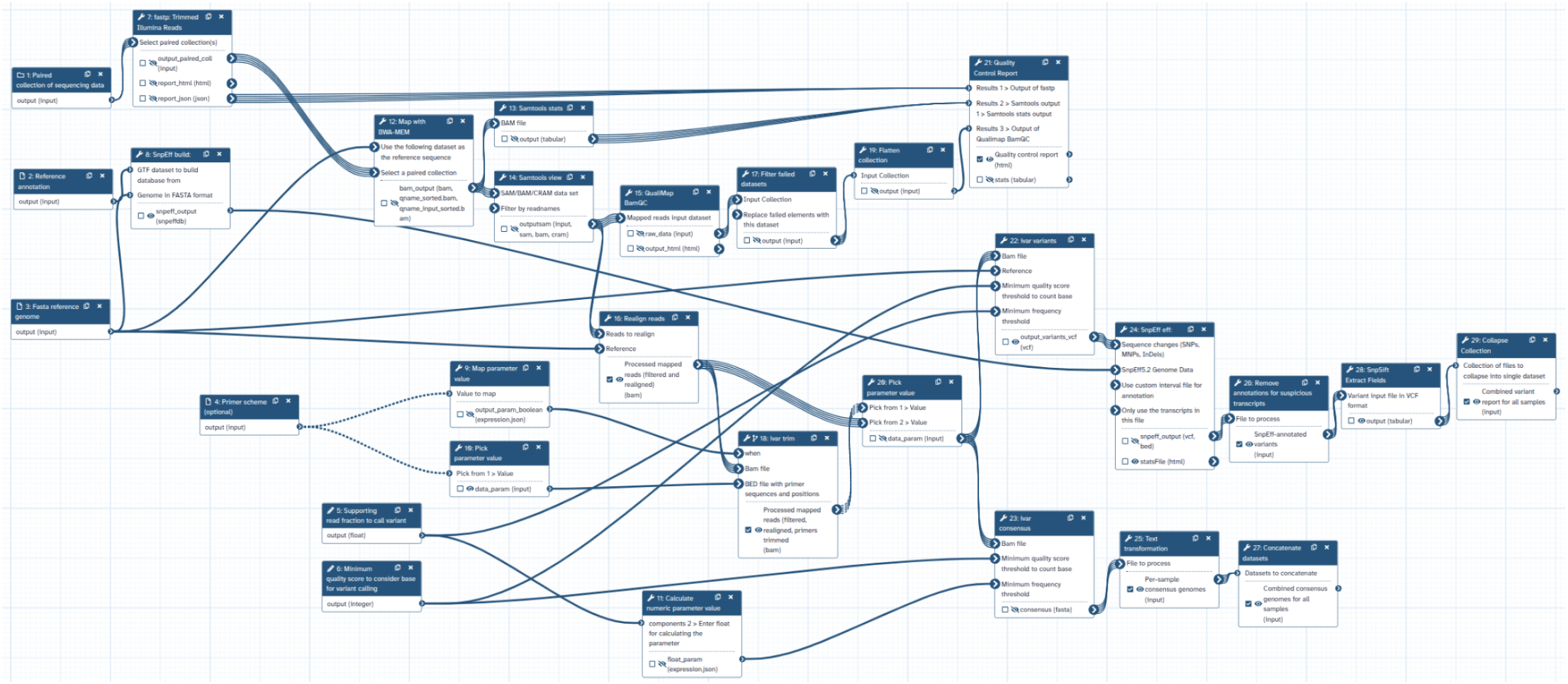
Schematics of the variant calling workflow used in this study (the workflow is available from [IWC](https://iwc.galaxyproject.org/workflow/generic-non-segmented-viral-variant-calling-main/)).

For a detailed workflow description see Methods. Briefly, reads are first quality- and adapter-trimmed with `fastp`, and primer sequences are removed with `iVar` trim when applicable. Cleaned reads are aligned to the reference using `BWA-MEM`, with BAM filtering, statistics, and alignment quality is assessed with `samtools` and `QualiMap BamQC`. Alignments are realigned around indels using the `LoFreq Viterbi` realigner to improve variant detection. Variants are then called with `iVar variants` under user-defined thresholds for base quality and allele frequency, and consensus genomes are generated with `iVar consensus`, introducing IUPAC ambiguity codes where mixed alleles occur. Variants are functionally annotated with `SnpEff` using a database built from the supplied reference, and results are parsed into tabular form with `SnpSift`. Quality metrics from all stages are summarized with `MultiQC`, while standard Galaxy collection and text utilities handle dataset organization and parameter mapping. The workflow outputs include trimmed reads, mapped and realigned BAMs, annotated VCFs with summary tables, per-sample consensus FASTA files, and integrated QC reports.

### Selecting SRA data and launching workflow (Video 1 & 2)

After selecting the reference genome and workflow for variant identification analysis, we chose public surveillance data for the virus. BRC-analytics’ SRA search wizard initially showed 466 individual datasets. These were automatically pre-filtered to 225 samples across nine studies, based on the workflow’s requirements for “paired end” (Illumina or Element) and “WGS” (whole genome sequencing) data; these filters can be adjusted as needed. Among these nine studies were the three largest measles surveillance datasets (Table 2), which we selected to launch the previously described workflow.

**Table 2.** Links and description for key artifacts of this study.

| Artifact | Link | Description |
| --- | --- | --- |
| Analysis history | https://usegalaxy.org/u/cartman/h/overlaps-in-measels-1 | Galaxy history is the workspace where all analyses are conducted. It stores all inputs, intermediate datasets and outputs. JupyterLite notebooks used in the analysis can also be found there. |
| Workflow | https://doi.org/10.5281/zenodo.16896089 | Variants calling workflow used to generate variant lists for this analysis. The history above contains intermediate and final data generated by this workflow. |
| Videos | https://gxy.io/brc-whitepaper-videos | Videos describing each stage of this study. |

**Table 2.**
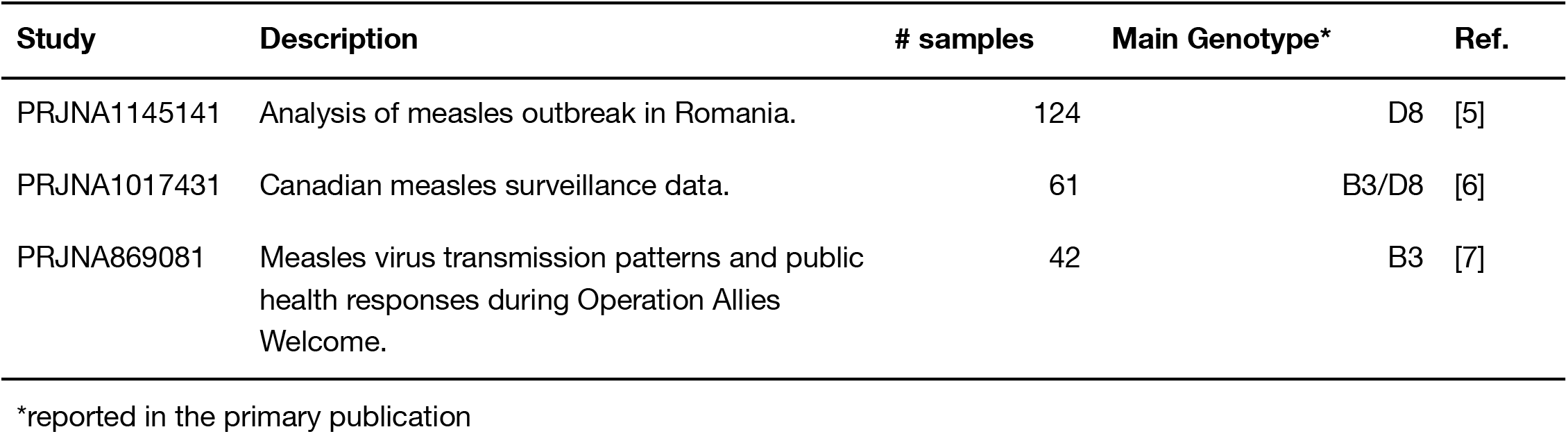
Three MeV surveillance datasets selected for this study.

| Study | Description | # samples | Main Genotype* | Ref. |
| --- | --- | --- | --- | --- |
| PRJNA1145141 | Analysis of measles outbreak in Romania. | 124 | D8 | [5] |
| PRJNA1017431 | Canadian measles surveillance data. | 61 | B3/D8 | [6] |
| PRJNA869081 | Measles virus transmission patterns and public health responses during Operation Allies Welcome. | 42 | B3 | [7] |

\*reported in the primary publication

The workflow outputs two datasets that are relevant to the evolution of overlapping coding regions in MeV. The first dataset contains information about each variant within each sample: its position in the genome, alternative allele frequency, strand-specific read counts, functional impact and other data. The second dataset reports consensus sequences produced for each sample by incorporating all supported variants present in that sample. We will use it to investigate adaptive significance of substitutions.

### Making sense of variants: Combining LLMs and JupyterLite (Video 3)

The workflow described above produced a list of variants observed in each sample. Such a list by itself does not provide any valuable biological insights. It needs to be reshaped, summarized, and visualized to give us any ideas on the biological significance of variants we just identified. Additional complication is that there are no “off-the-shelf” tools for analysis of genetic variants in overlapping coding regions. BRC-analytics and Galaxy (Fig. 5) provide an ability to construct analyses in an *ad-hoc* way, using an environment called JupyterLite. The entire analysis including the notebook can be accessed as described in Methods (also see the Video 3). All steps of this analysis described in the notebook were generated with the help of ChatGPT using a setup shown in Fig. 5 in which we used two open browser windows: one with Galaxy workspace and another with ChatGPT, Microsoft CoPilot, Claude, or any other LLM agent capable of writing Python code (the video also details safety checks we used when using LLM-generated code—such code should ALWAYS be treated with caution and made publicly available).

**Figure 5.**
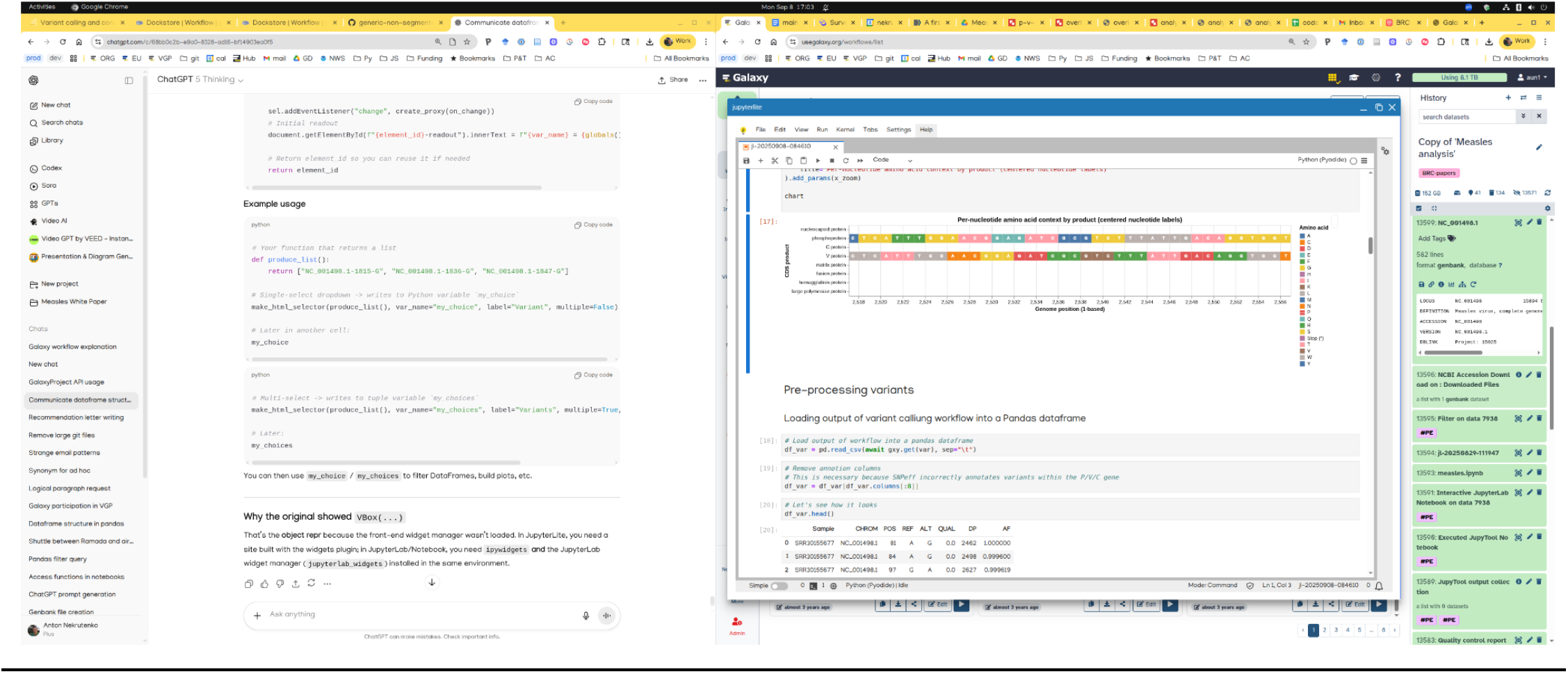
Analysis setup for post-processing of variant calls.

### P reading frame is more permissive than C and V (Video 3)

First (Fig. 6) we looked for general patterns of substitutions within P/C and P/V overlaps by simply plotting aggregate data for each substitution (e.g., the number of samples a substitution occurs at, median, min, and max alternative allele frequencies and its effect, synonymous or non-synonymous, on each reading frame). Visually, the P reading frame appears to be more permissive or genetically plastic —most substitutions are non-synonymous (“red”) in P and synonymous (“green”) in C or V.

**Figure 6.**
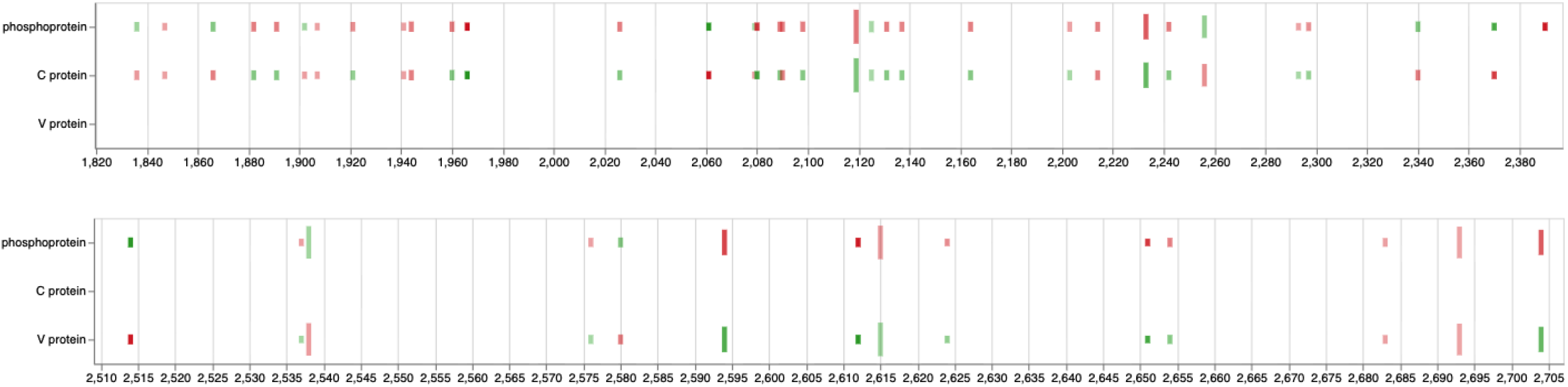
Nucleotide changes within P/C (top) and P/V (bottom) overlaps. Each substitution is represented by two ticks corresponding to each reading frame. Green = synonymous for that frame; Red = non-synonymous for that frame; length of each tick represents the alternative allele frequency spread = the difference between min and max values. The opacity is the number of samples (from min = 2 to max = 225) the change is found in. A live version of this notebook can be viewed [here](https://usegalaxy.org/visualizations/display?visualization=jupyterlite&dataset_id=f9cad7b01a47213516812bfc3ef85974).

Nucleotide substitutions in the C-protein and the V-protein region downstream of the editing site appear to affect C and V protein function less than they affect protein P (Table 3). Variants in the C–P overlap (nucleotides ∼1,836–2,370) are mostly synonymous in C but nonsynonymous in P. This suggests strong purifying selection on C (which contains nuclear localization, export, and SHCBP1-binding signals), while P tolerates or favors amino-acid changes in this region, indicating adaptive fine-tuning of P’s polymerase cofactor and regulatory functions while C’s integrity is maintained. Substitutions in the P–V overlap (nucleotides ∼2,514–2,704) frequently show synonymous changes in V but nonsynonymous in P, again pointing to P-directed adaptation. However, some sites (e.g., 2514-A, 2580-G) show the opposite pattern (nonsynonymous in V, synonymous in P), possibly corresponding to residues within or adjacent to V/P amino-acid positions 100–120, a region known to mediate STAT1 and Jak1 binding [8,9]. These variants could represent V-specific adaptive evolution affecting interferon antagonism, while the broader pattern reflects the evolutionary compromise needed to maintain two overlapping reading frames with distinct but interdependent functions (see more on this in the evolutionary analysis section below).

**Table 3.**
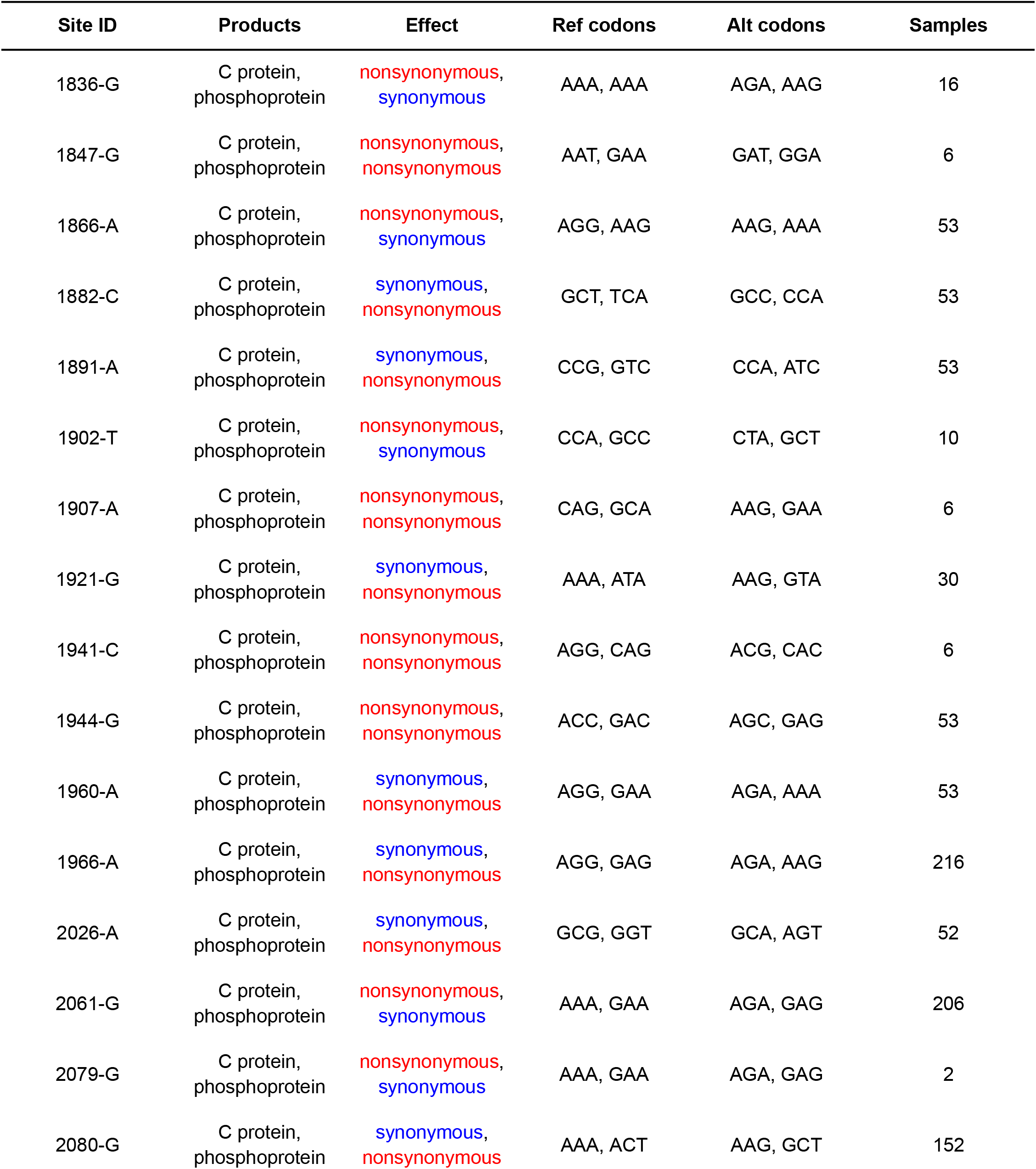
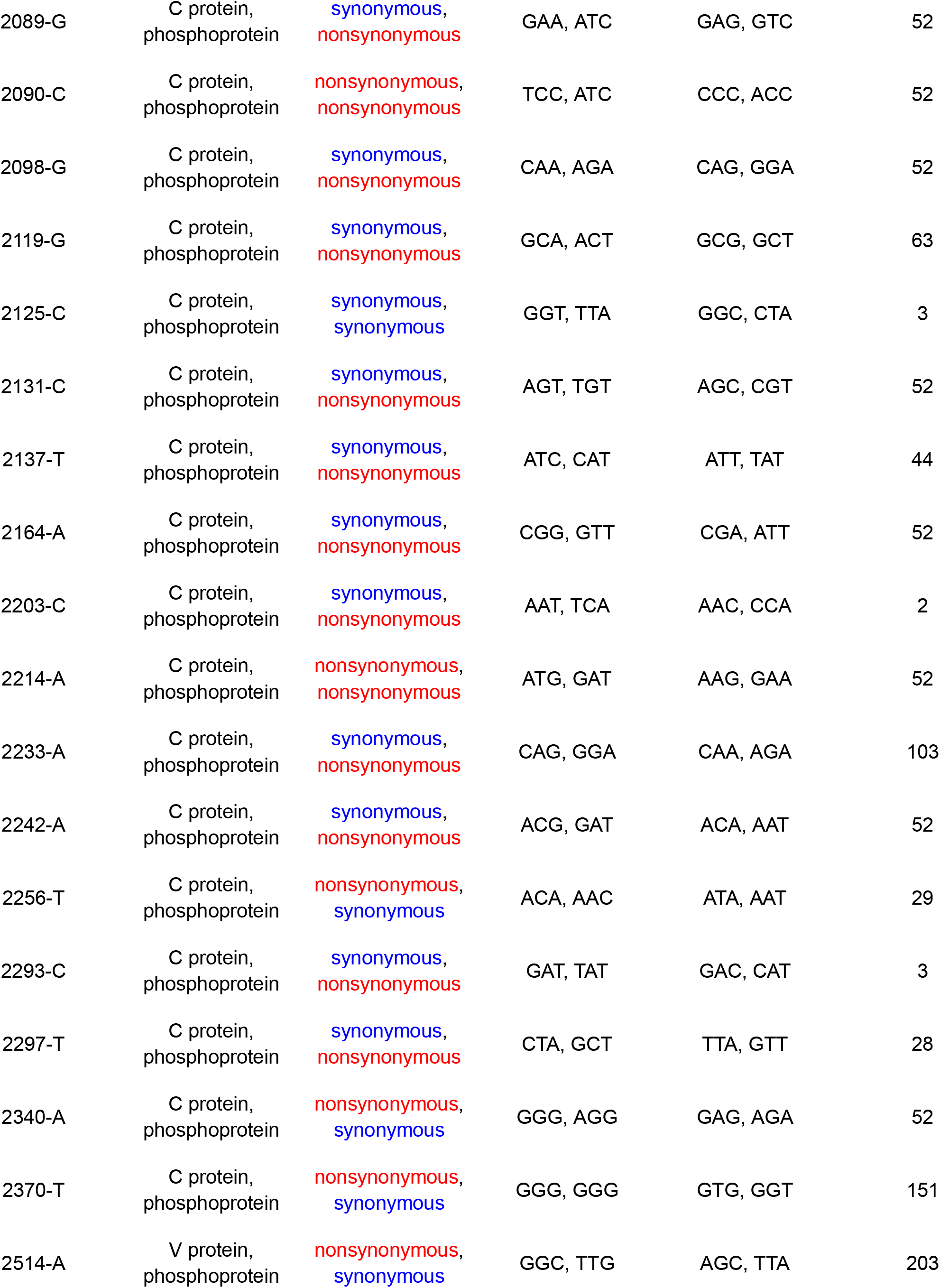
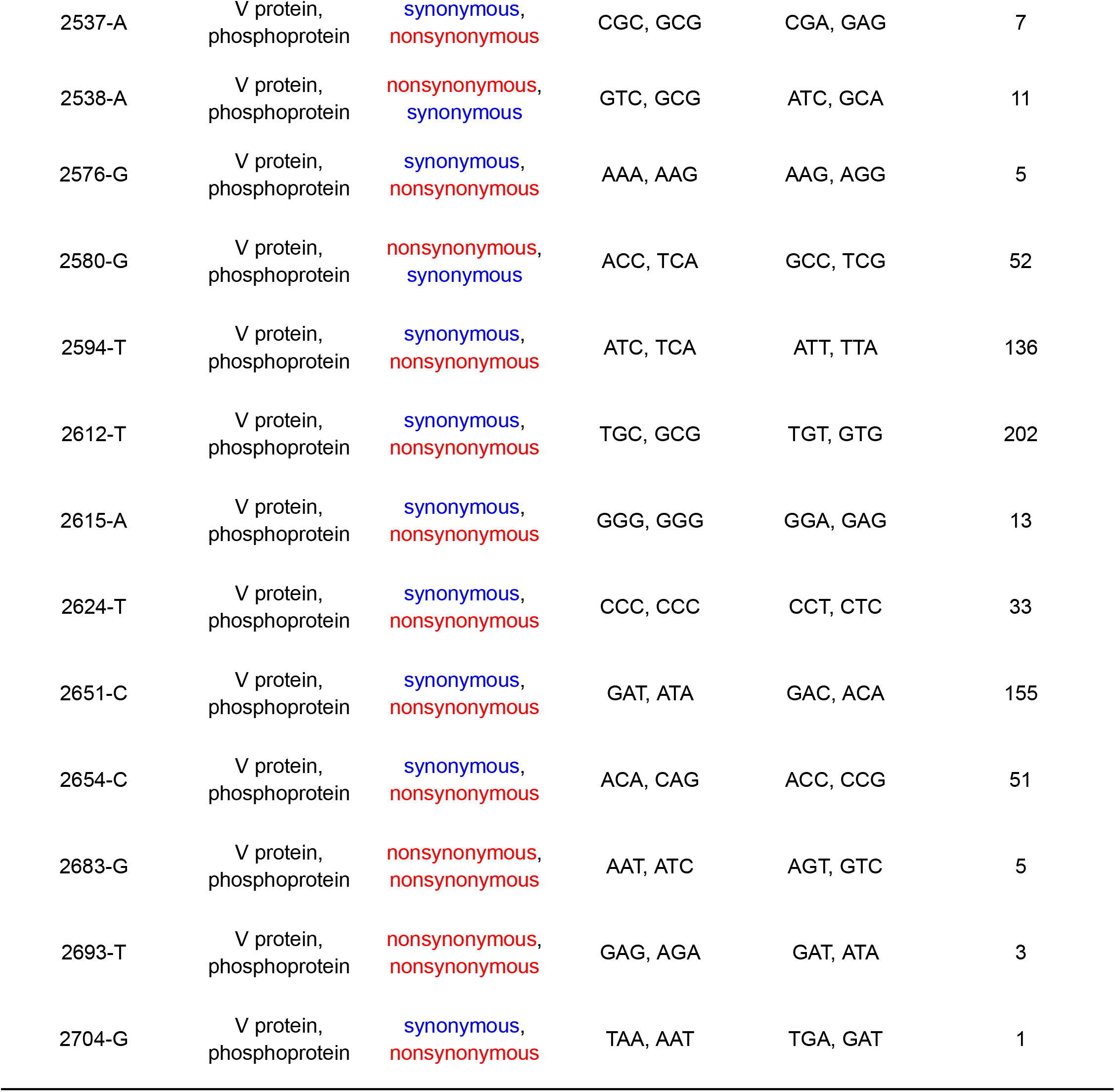
Nucleotide substitutions within P/C and P/V overlaps. Samples = number of samples this variant is observed in.

| Site ID | Products | Effect | Ref codons | Alt codons | Samples |
| --- | --- | --- | --- | --- | --- |
| 1836-G | C protein, phosphoprotein | nonsynonymous, synonymous | AAA, AAA | AGA, AAG | 16 |
| 1847-G | C protein, phosphoprotein | nonsynonymous, nonsynonymous | AAT, GAA | GAT, GGA | 6 |
| 1866-A | C protein, phosphoprotein | nonsynonymous, synonymous | AGG, AAG | AAG, AAA | 53 |
| 1882-C | C protein, phosphoprotein | synonymous, nonsynonymous | GCT, TCA | GCC, CCA | 53 |
| 1891-A | C protein, phosphoprotein | synonymous, nonsynonymous | CCG, GTC | CCA, ATC | 53 |
| 1902-T | C protein, phosphoprotein | nonsynonymous, synonymous | CCA, GCC | CTA, GCT | 10 |
| 1907-A | C protein, phosphoprotein | nonsynonymous, nonsynonymous | CAG, GCA | AAG, GAA | 6 |
| 1921-G | C protein, phosphoprotein | synonymous, nonsynonymous | AAA, ATA | AAG, GTA | 30 |
| 1941-C | C protein, phosphoprotein | nonsynonymous, nonsynonymous | AGG, CAG | ACG, CAC | 6 |
| 1944-G | C protein, phosphoprotein | nonsynonymous, nonsynonymous | ACC, GAC | AGC, GAG | 53 |
| 1960-A | C protein, phosphoprotein | synonymous, nonsynonymous | AGG, GAA | AGA, AAA | 53 |
| 1966-A | C protein, phosphoprotein | synonymous, nonsynonymous | AGG, GAG | AGA, AAG | 216 |
| 2026-A | C protein, phosphoprotein | synonymous, nonsynonymous | GCG, GGT | GCA, AGT | 52 |
| 2061-G | C protein, phosphoprotein | nonsynonymous, synonymous | AAA, GAA | AGA, GAG | 206 |
| 2079-G | C protein, phosphoprotein | nonsynonymous, synonymous | AAA, GAA | AGA, GAG | 2 |
| 2080-G | C protein, phosphoprotein | synonymous, nonsynonymous | AAA, ACT | AAG, GCT | 152 |
| 2089-G | C protein,<br>phosphoprotein | synonymous,<br>nonsynonymous | GAA, ATC | GAG, GTC | 52 |
| 2090-C | C protein,<br>phosphoprotein | nonsynonymous,<br>nonsynonymous | TCC, ATC | CCC, ACC | 52 |
| 2098-G | C protein,<br>phosphoprotein | synonymous,<br>nonsynonymous | CAA, AGA | CAG, GGA | 52 |
| 2119-G | C protein,<br>phosphoprotein | synonymous,<br>nonsynonymous | GCA, ACT | GCG, GCT | 63 |
| 2125-C | C protein,<br>phosphoprotein | synonymous,<br>synonymous | GGT, TTA | GGC, CTA | 3 |
| 2131-C | C protein,<br>phosphoprotein | synonymous,<br>nonsynonymous | AGT, TGT | AGC, CGT | 52 |
| 2137-T | C protein,<br>phosphoprotein | synonymous,<br>nonsynonymous | ATC, CAT | ATT, TAT | 44 |
| 2164-A | C protein,<br>phosphoprotein | synonymous,<br>nonsynonymous | CGG, GTT | CGA, ATT | 52 |
| 2203-C | C protein,<br>phosphoprotein | synonymous,<br>nonsynonymous | AAT, TCA | AAC, CCA | 2 |
| 2214-A | C protein,<br>phosphoprotein | nonsynonymous,<br>nonsynonymous | ATG, GAT | AAG, GAA | 52 |
| 2233-A | C protein,<br>phosphoprotein | synonymous,<br>nonsynonymous | CAG, GGA | CAA, AGA | 103 |
| 2242-A | C protein,<br>phosphoprotein | synonymous,<br>nonsynonymous | ACG, GAT | ACA, AAT | 52 |
| 2256-T | C protein,<br>phosphoprotein | nonsynonymous,<br>synonymous | ACA, AAC | ATA, AAT | 29 |
| 2293-C | C protein,<br>phosphoprotein | synonymous,<br>nonsynonymous | GAT, TAT | GAC, CAT | 3 |
| 2297-T | C protein,<br>phosphoprotein | synonymous,<br>nonsynonymous | CTA, GCT | TTA, GTT | 28 |
| 2340-A | C protein,<br>phosphoprotein | nonsynonymous,<br>synonymous | GGG, AGG | GAG, AGA | 52 |
| 2370-T | C protein,<br>phosphoprotein | nonsynonymous,<br>synonymous | GGG, GGG | GTG, GGT | 151 |
| 2514-A | V protein,<br>phosphoprotein | nonsynonymous,<br>synonymous | GGC, TTG | AGC, TTA | 203 |
| 2537-A | V protein,<br>phosphoprotein | synonymous,<br>nonsynonymous | CGC, GCG | CGA, GAG | 7 |
| 2538-A | V protein,<br>phosphoprotein | nonsynonymous,<br>synonymous | GTC, GCG | ATC, GCA | 11 |
| 2576-G | V protein,<br>phosphoprotein | synonymous,<br>nonsynonymous | AAA, AAG | AAG, AGG | 5 |
| 2580-G | V protein,<br>phosphoprotein | nonsynonymous,<br>synonymous | ACC, TCA | GCC, TCG | 52 |
| 2594-T | V protein,<br>phosphoprotein | synonymous,<br>nonsynonymous | ATC, TCA | ATT, TTA | 136 |
| 2612-T | V protein,<br>phosphoprotein | synonymous,<br>nonsynonymous | TGC, GCG | TGT, GTG | 202 |
| 2615-A | V protein,<br>phosphoprotein | synonymous,<br>nonsynonymous | GGG, GGG | GGA, GAG | 13 |
| 2624-T | V protein,<br>phosphoprotein | synonymous,<br>nonsynonymous | CCC, CCC | CCT, CTC | 33 |
| 2651-C | V protein,<br>phosphoprotein | synonymous,<br>nonsynonymous | GAT, ATA | GAC, ACA | 155 |
| 2654-C | V protein,<br>phosphoprotein | synonymous,<br>nonsynonymous | ACA, CAG | ACC, CCG | 51 |
| 2683-G | V protein,<br>phosphoprotein | nonsynonymous,<br>nonsynonymous | AAT, ATC | AGT, GTC | 5 |
| 2693-T | V protein,<br>phosphoprotein | nonsynonymous,<br>nonsynonymous | GAG, AGA | GAT, ATA | 3 |
| 2704-G | V protein,<br>phosphoprotein | synonymous,<br>nonsynonymous | TAA, AAT | TGA, GAT | 1 |

### Substitutions have distinct geographic and temporal patterns (Video 3)

Next we analyzed the substitution patterns in the context of sampling locations and collection times. For this we had to rely on the sample metadata downloaded from SRA. Results of this analysis are shown in Fig. 7. While most samples show strain-specific patterns there is a number if interesting sites such as, for example linked co-occurring substitutions at position 2,576 and 2,683 within five US samples (SRR25426258, SRR25426259, SRR25426260, SRR25426262, SRR25426264). These two sites are within the P/V overlap.

**Figure 7.**
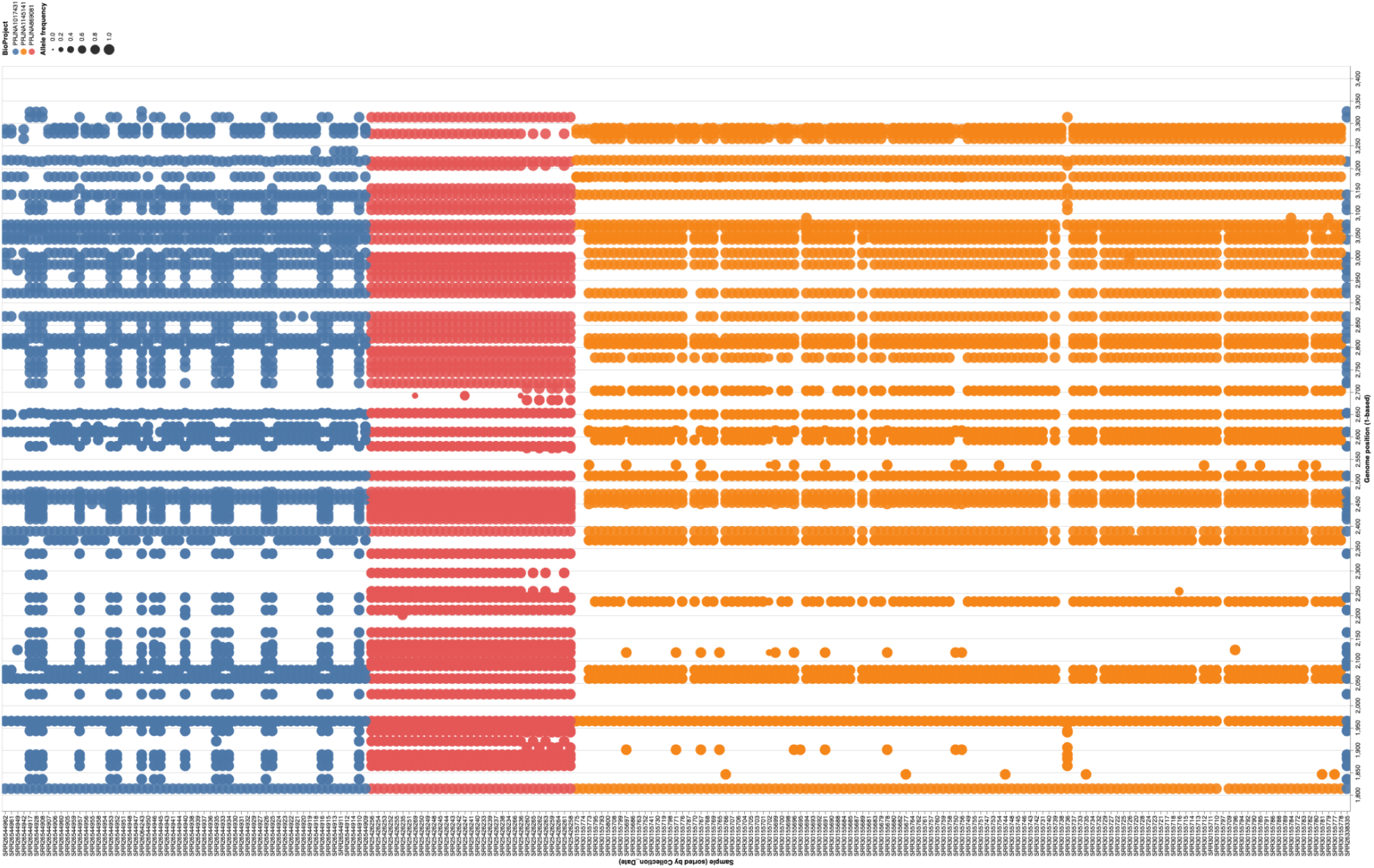
Substitutions within P/V/C locus in each sample. Samples (X-axis) are sorted by collection time from 01/22/2018 (left) to 05/07/2024 (right). The rightmost blue sample had no collection date associated with it. Genomic coordinates of variants are on the Y-axis. Blue = Canada; Red = USA; Orange = Romania. Size of this circle is proportional for alternative allele frequency at that site. A live version of this notebook can be viewed [here](https://usegalaxy.org/visualizations/display?visualization=jupyterlite&dataset_id=f9cad7b01a47213516812bfc3ef85974).

The first change is non-synonomous in P and synonymous in V, while the second is non-synonymous for both reading frames.

### Comparative analysis of substitution effects (Video 4)

Our final analysis aimed to identify potentially adaptive substitutions, particularly within the P/C and P/V overlaps. Traditional codon-based models, which assume a constant nonsynonymous to synonymous substitution rate (ω = dN/dS) across all lineages at a given site, are not well-suited for detecting transient or lineage-specific instances of positive selection. To address this limitation, we used the Mixed Effects Model of Evolution (MEME) method [10]. MEME is specifically designed to detect episodic diversifying selection at individual sites within a multiple sequence alignment. It achieves this by integrating fixed effects (allowing ωto vary across sites) with random effects (allowing ω to vary across branches). For each site, MEME fits a mixture model. This model accounts for the possibility that some branches may be subject to purifying or neutral selection (ω ≤ 1), while others may experience positive selection (ω > 1). The probabilities of these scenarios are estimated using maximum likelihood. This approach enables MEME to pinpoint sites where positive selection has occurred in only a subset of lineages, rather than uniformly across the entire phylogeny.

To apply MEME directly to the output of our workflow we will use consensus sequences generated by the variant caller for each processed sample. These sequences can be used as an input to the MEME tool that already exists in the Galaxy platform. However, MEME requires exact codon alignments of the coding sequences to perform the detection of sites under selection. Thus before we can use MEME we need to post-process sequences generated by the workflow. Specifically, we need extract regions corresponding to P, V, and C proteins. For this we would use a new JupyterLite notebook ([Video 4](https://youtu.be/opPVkPxU25c)). Similarly to the previous analysis this notebook was generated entirely using ChatGPT LLM. It takes two files from the Galaxy history: the GenBank annotation for measles and consensus sequences generated by the variant calling workflow.

The notebook first filters out sequences that differ in length from the MeV genome. Next, it reads the coordinates of CDS annotations from the GenBank file and uses these coordinates to slice consensus sequences into CDS blocks, serving each separately. We then save consensus sequences corresponding to CDS for P, V, and C proteins back to the Galaxy, use Rapid Neighbour-Joining tool [11] in Galaxy to compute phylogenetic trees for each set of sequences and obtain MEME estimates. To eliminate potential artifacts this analysis was performed exclusively using interior branches of the tree [12].

Two sites stand out as potentially showing evidence of episodic diversifying positive selection: codons 105 and 111 of the P/V proteins (corresponding to codons 97 and 103 of the C proteins; Fig. 8). These are located before the RNA editing site within the P/V reading frame and have identical effects on the P and V products. Both substitutions are non-synonymous on the P/V frame and synonymous in the C-frame.

**Figure 8.**
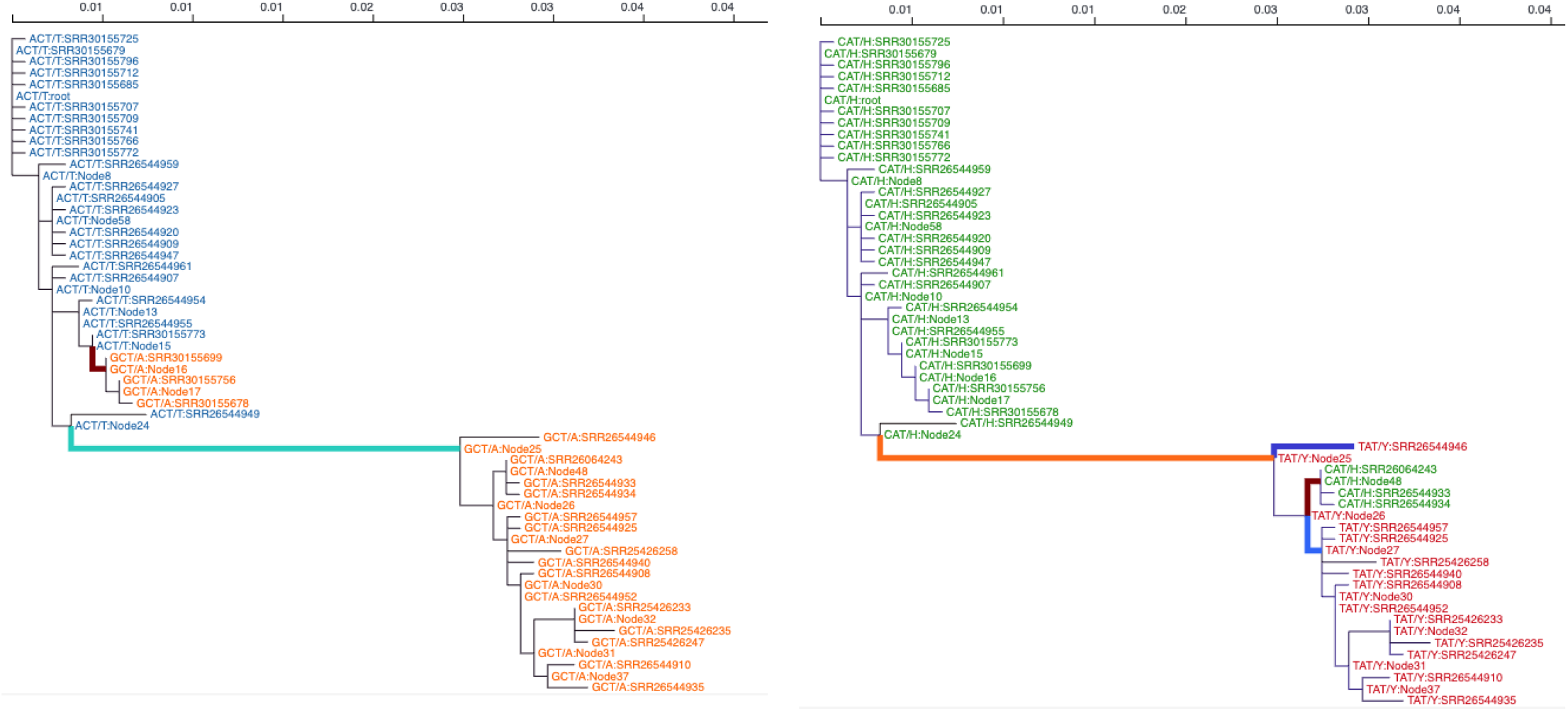
Tracing substitutions within codons 105 (left) and 111 (right) of the P/V reading frames through the phylogenetic tree (unrooted) of the analyzed samples. Imputed states at internal nodes are also shown. The number of samples shown here is smaller than the total number of samples in our analysis because identical sequences were excluded from the analysis. Sequences with an excessive number of unresolved (N) characters were also excluded.

Both sites are within the V/P “anti-STAT1” hot zone (codons 110 - 115; [8]). Residue 105 lies upstream of that motif in the same PNT (P-N-terminal) domain that mediates interactions with STAT1/Jak1. Changes here could allosterically affect the 110–120 epitope (local secondary structure/loop positioning) and thus tune antagonism strength—similar in principle to the well-characterized Y110H loss-of-function effect [13][14].

Residue 111 is embedded in the P/V common N-terminal STAT1-binding segment (∼110–120 aa). This segment contains the proven triad Y110, V112, H115—mutating any of those disrupts STAT1 antagonism and attenuates virus in vivo. Variation at 111 is immediately adjacent to V112 and within the same interface; it could modulate binding geometry/affinity to STAT1 (and also Jak1), shifting the potency of IFN-α/β and IFN-γ blockade [9,15,16]. In addition, position 111 is located within an experimentally confirmed T-cell epitope [17], where the Y to H mutation (found in the D8 MeV genotype) reduces the potency of T-cell response compared to the wildtype (and the vaccine strain).

Two residues under episodic diversifying selection (positions 105 and 111) fall within the shared N-terminal region of the measles virus V protein, adjacent to the experimentally validated STAT1-binding triad (Y110, V112, H115 [8]). Given the essential role of this motif in interferon antagonism, variation at 105 and 111 could adaptive modulation of host–virus interactions rather than neutral drift. These sites may fine-tune V’s capacity to sequester STAT1 or balance replication efficiency with innate immune evasion, consistent with prior observations that alterations in this region modulate viral attenuation and immune suppression.

## Summary

BRC-Analytics offers a complete, browser-based solution for reproducible genomic analysis of infectious diseases. In this example analysis, we showed how it allows users to select the *Morbillivirus hominis* reference genome, apply community-curated workflows for variant calling and query NCBI SRA for 225 surveillance samples. The platform integrates `JupyterLite` notebooks, assisted by large language models (LLMs), to facilitate rapid visualization and interpretation of substitution patterns. This reveals distinct geographic and temporal clustering, and shows a lesser functional impact for C and V protein substitutions compared to P protein. Comparative evolutionary analysis using the Mixed Effects Model of Evolution (MEME) method, identified two sites (codons 105 and 111) in the V/P overlap region that are under episodic diversifying selection. These sites are adjacent to the STAT1-binding motif, suggesting an adaptive role in host-virus interactions, and have also been characterized as belonging to a CTL-epitope with one mutation (Y111H) showing an experimental reduction in T-cell response.

By unifying reference selection, workflow execution, data retrieval, and post-analysis within a single browser session, BRC-Analytics eliminates the need for local installations and manual data transfers, ensuring reproducible and interactive evolutionary and functional analyses.

## Methods

### Viral variant calling and consensus construction

All analyses were performed within the Galaxy platform (https://usegalaxy.org) using the publicly available release 0.1 of the workflow “*Variant calling and consensus construction from paired-end short-read data of non-segmented viral genomes*”. This workflow automates read quality control, alignment, variant detection, consensus genome reconstruction, and annotation in a fully reproducible environment.

Paired-end Illumina reads were first quality-filtered and adapter-trimmed with `fastp` v1.0.1 [18] using default parameters. When an amplicon primer scheme was provided, primer sequences were removed with `iVar trim` v1.4.4 [19]. Filtered reads were aligned to the appropriate reference genome with `BWA-MEM` v0.7.19 [20], and alignment quality was assessed using `samtools stats` v2.0.7 and `QualiMap BamQC` v2.3 [21].

To improve indel representation before variant calling, alignments were realigned with the `LoFreq` Viterbi realigner v2.1.5 [22]. Variants were then identified using `iVar variants` v1.4.4, requiring a minimum base quality of 20 and an alternate-allele frequency ≥ 0.25 by default. Per-sample consensus genomes were generated with `iVar consensus` v1.4.4, inserting IUPAC ambiguity codes at positions where minor variants exceeded the user-defined threshold (typically 25%).

Functional annotation of single-nucleotide variants was performed using `SnpEff` v5.2 [23], with a local database built directly from the supplied reference GenBank file (Note that these were not used in the analysis. Instead annotation was re-done in JupyterLite to account for overlapping reading frames). Annotated variants were extracted into tabular form via `SnpSift ExtractFields` v5.2 [24]. Summary quality metrics from `fastp`, `samtools`, and `QualiMap` were collated into a unified interactive report using `MultiQC` v1.24.1 [25].

All dataset handling (collection flattening, parameter mapping, concatenation, and filtering) was executed using standard Galaxy collection and text-manipulation utilities, ensuring full provenance capture. The workflow yields cleaned reads, alignment files (BAM), variant calls (VCF), annotated variant tables, consensus FASTA sequences, and a comprehensive MultiQC report for each batch of samples.

## Acknowledgements

We also would like to express our immense gratitude to Dan Stanzione and David Hancock for essential computational resources provided by the Advanced Cyberinfrastructure Coordination Ecosystem (ACCESS-CI), Texas Advanced Computing Center, and the JetStream2 scientific cloud. This work is funded by the NIH Grant U24AI183870.

